# NK cells negatively regulate CD8 T cells to promote immune exhaustion and chronic *Toxoplasma gondii* infection

**DOI:** 10.1101/864272

**Authors:** Daria L. Ivanova, Ryan Krempels, Stephen L. Denton, Kevin D. Fettel, Giandor M. Saltz, David Rach, Rida Fatima, Tiffany Mundhenke, Joshua Materi, Ildiko R. Dunay, Jason P. Gigley

## Abstract

NK cells regulate CD4+ and CD8+ T cells in acute viral infection, vaccination and the tumor microenvironment. NK cells also become exhausted in chronic activation settings. The mechanisms causing these ILC responses and their impact on adaptive immunity are unclear. CD8+ T cell exhaustion develops during chronic *Toxoplasma gondii* (*T. gondii*) infection resulting in parasite reactivation and death. How chronic *T. gondii* infection impacts the NK cell compartment is not known. We demonstrate that NK cells do not exhibit hallmarks of exhaustion. Their numbers are stable and they do not express high PD1 or LAG3. NK cell depletion with anti-NK1.1 is therapeutic and rescues chronic *T. gondii* infected mice from CD8+ T cell exhaustion dependent death, increases survival after lethal secondary challenge and reduces parasite reactivation. Anti-NK1.1 treatment increased polyfunctional CD8+ T cell responses in spleen and brain and reduced CD8+ T cell apoptosis. Chronic *T. gondii* infection promotes the development of a modified NK cell compartment, which does not exhibit normal NK cell behavior. This splenic CD49a-CD49b+NKp46+ NK cell population develops during the early chronic phase of infection and increases through the late chronic phase of infection. They are Ly49 and TRAIL negative and are enriched for expression of CD94/NKG2A and KLRG1. They do not produce IFNγ, are IL-10 negative, do not increase PDL1 expression, but do increase CD107a on their surface. They are also absent from brain. Based on the NK cell receptor phenotype we observed NKp46 and CD94-NKG2A cognate ligands were measured. Activating NKp46 (NCR1-ligand) ligand increased and NKG2A ligand Qa-1b expression was reduced. Blockade of NKp46 also rescued the chronically infected mice from death. Immunization with a single dose non-persistent 100% protective *T. gondii* vaccination did not induce this cell population in the spleen, suggesting persistent infection is essential for their development. We hypothesize chronic *T. gondii* infection induces an NKp46 dependent modified NK cell population that reduces functional CD8+ T cells to promote persistent parasite infection in the brain. NK cell targeted therapies could enhance immunity in people with chronic infections, chronic inflammation and cancer.

## Introduction

*Toxoplasma gondii* (*T. gondii*) is an obligate intracellular protozoan that is the 3^rd^ leading cause of foodborne illness in the U.S.[1] At least one-third of the human population is infected with this parasite and it is a major health concern for people who become immune compromised and in the developing fetus[2; 3]. Presently, there are no vaccines or drugs available to prevent or eliminate this infection and infection with this parasite is life long [4; 5]. *T. gondii* infection induces a potent cell mediated response that is initiated by the production of IL-12 which helps activate CD8+ T cells to produce IFNγ[6; 7; 8; 9]. CD8+ T cell IFNγ production is the major mediator of this infection. Despite induction of a robust Th1 response, the parasite is never cleared. The immunological reason why this infection is not cleared is still unknown.

In mouse models of chronic *T. gondii* infection the parasite can spontaneously reactivate causing the development of toxoplasmic encephalitis (TE) and death [10]. Parasite reactivation has been attributed to the development of immune exhaustion of parasite specific CD8+ T cells[10; 11; 12; 13]. The CD8+ T cells in mice harboring chronic *T. gondii* infection exhibit immune exhaustion characteristics similar to persistent viral infections[14]. Loss of activated CD8+ T cells resulting in a reduced functional cell population, expression of high levels of programmed death 1(PD1) and increased apoptosis of CD8+ T cells. This loss of functional CD8+ T cells results in parasite reactivation and death of the animals. Importantly, the exhausted CD8+ T cells can be rescued with anti-PDL1 therapy during chronic *T. gondii* infection and this also prevents parasite reactivation and death. The mechanisms underlying the development of CD8+ T cell exhaustion and dysfunction during chronic *T. gondii* infection are still unclear.

NK cells are innate lymphoid cells (ILCs) that provide early cytotoxicity and cytokine dependent protection during infections and cancer [15]. NK cells are important for control of acute *T. gondii* infection[16; 17] and are activated early during parasite infection by IL-12[18; 19]. As a result of IL-12 signaling, NK cells produce high levels of IFNγ, which helps control the parasite prior to T cell activation. NK cells are more complex than previously thought and appear to not only be activated and work as a component of innate immunity during acute infections, but may also continue to work along side CD4+ and CD8+ T cells during the adaptive phase of immunity. NK cells have been shown to acquire memory-like features after exposure to haptens, during viral infections and after cytokine stimulation [20; 21; 22; 23]. This highlights their ability to not simply fall into the background once adaptive immunity is established, but also to continue to play a role in immunity after acute infections are resolved. NK cells have also been shown to become exhausted[24; 25; 26; 27]. This can occur in the tumor microenvironment, chronic stimulation and persistent HCV infection. In these different disease situations, NK cells become dysfunctional and as a result could contribute to the persistence of infections and reduced clearance of tumor cells. NK cells can alsp be negative regulators of the adaptive response during acute infections and cancer. Through several interactions including TRAIL, NKp46 and yet to be defined receptors, NK cells can lyse CD4+ and CD8+ T cells resulting in less effective adaptive responses thereby promoting pathogen and tumor persistence[28; 29; 30; 31; 32; 33]. In addition, NK cells produce IL-10 during acute systemic infections including *T. gondii* infection dampening the activation of adaptive immune responses[34]. Much of what is known about the development of these other non-protective NK cell responses is in the acute disease or infection setting and less in known about how NK cells behave during chronic infections long after acute infection is resolved.

Based upon the knowledge that CD8+ T cells become exhausted to promote *T. gondii* persistence, NK cells can remain active for long periods of time, NK cells have the potential to become exhausted and they can regulate development of adaptive immune responses we were interested to test how chronic *T. gondii* infection impacted the NK cells and how did NK cells impact the outcomes of chronic toxoplasmosis. Our results indicate that NK cells are still present during chronic *T. gondii* infection. They do not exhibit characteristics of immune exhaustion. They contribute to the loss of exhausted CD8+ T cells and their removal helps maintain control of chronic *T. gondii* infection. We also demonstrate that NK cells develop a unique phenotype that supports the hypothesis that NKp46 recognition of ligand and loss of NKG2A interaction with Qa-1b promotes the development of an NK cell population that negatively regulates CD8+ T cell function resulting in parasite reactivation and death. Our data highlight that NK cells could be therapeutic targets to enhance long-term immunity to chronic *T. gondii* infection.

## Materials and methods

### Mice

C57BL/6 (B6), B6.129S6-IL-10^tm1Flv^/J (IL-10-GFP Tiger) mice were purchased from The Jackson Laboratory. All animals were housed under specific pathogen-free conditions at the University of Wyoming Animal Facility. This study was carried out in strict accordance following the recommendations in the Guide for the Care and Use of Laboratory Animals of the National Institutes of Health. The University of Wyoming Institutional Animal Care and Use Committee (IACUC) (PHS/NIH/OLAW assurance number: A3216-01) approved all animal protocols.

### T. gondii *parasites and infection*

Tachyzoites of RH were cultured by serial passage in human fetal lung fibroblast (MRC5, ATCC) cell monolayers in complete DMEM (supplemented with 0.2 mM uracil for CPS strain). For mouse infections, parasites were purified by filtration through a 3.0-µm filter (Merck Millipore Ltd.) and washed with phosphate-buffered saline (PBS). Mice were infected intraperitoneally (i.p.) with 1 × 10^3^ or 1 × 10^6^ RH tachyzoites or 1 × 10^6^ CPS tachyzoites. The brains of CBA mice 5 weeks after ME49 infection were used as a source of ME49 cysts. Mice were infected i.p. or i.g. (intragastrically) with 10 or 200 ME49 cysts.

### NK Cell depletion and Nkp46 blockade in vivo

To deplete NK cells, B6 mice were treated i.p. with 200 μg of anti-NK1.1 (PK136, Bio X Cell). To block NKp46 mice were treated i.p. with 50 ug non-depleting LEAF purified anti-NKp46 (29A1.4, Biolegend)[35]. Antibody treatments were started 5 weeks after infection with ME49 and continued every other day for 2 weeks for flow cytometry assays or until non-treated animal groups died from reactivation of *T. gondii*.

### Brain and spleen T cell isolation and stimulation

Single-cell suspensions of brain and spleen were prepared from mice. To harvest brain lymphocytes and assess their phenotype and function, mice were anesthetized and perfused with 20 mls of 0.9% saline with heparin as described[36]. Brains were then homogenized in 1 X PBS using a dounce homogenizer. Brains were pelleted by centrifugation then homogenates were added to 30% percoll® and centrifuged at 2000xg for 20 minutes at 15°C to collect lymphocytes from the pellet. Brain lymphocytes were then plated at 0.5 `−`1.5 × 10^6^ cells/well in complete Iscove’s DMEM medium (10% FBS, Na Pyruvate, non essential amino acids, penicillin, β-2 Mercaptoethanol) (Corning). 0.5 × 10^6^ congenically marked CD45.1 splenocytes were also added to the brain lymphocyte wells as feeder cells for antigen restimulation. Spleens were crushed through 70 um cell strainers (VWR) in 1 X PBS. Splenocytes were then treated with 3 ml of RBC lysis buffer for 3 minutes at 37C to lyse erythrocytes, washed then resuspended in complete Iscoves DMEM. Spleen cells were plated at 1 × 10^6^ cells per well. Brain and spleen cells were then pulsed with 20ug/ml Toxoplasma lysate antigen (TLA) for 8 hours and cultured at 37C in 5% CO2. After 8 hours, 1× protein transport inhibitor cocktail (PTIC) containing Brefeldin A/Monensin (eBioscience, Thermo Fisher Scientific) with or without anti-CD107a (eBio1D4B, eBioscience, Thermo Fisher Scientific) was added to each well in complete Iscove’s DMEM medium (Corning). After 4 hours incubation at 37C in 5% CO2, cells were prepared for flow cytometry.

### ILC functional assays

For ILC function assays, spleen cells were stimulated for 4 h with plate bound anti-NK1.1 in the presence of 1 × protein transport inhibitor cocktail (PTIC) containing Brefeldin A/Monensin (eBioscience, Thermo Fisher Scientific) and anti-CD107a (eBio1D4B, eBioscience, Thermo Fisher Scientific) in complete Iscove’s DMEM medium (Corning). Cells were incubated during stimulation at 37C in 5% CO2 for 4 hours. Cells were then first surface stained then intracellularly stained to measure function. ILC phenotypes were measured directly ex vivo. Spleen cells were stained following procedures indicated below after fixable Live/Dead staining (Invitrogen).

### Flow cytometry

Single cell suspensions from brain or spleen were assayed for immune cell phenotype and functions. Phenotype assays were performed directly ex vivo after harvest. Function assays were performed after antigen pulse cells or stimulation. All flow cytometry staining was performed using the same procedure for all experiments. Cells were washed twice with PBS and stained for viability in PBS using Fixable Live/Dead Aqua (Invitrogen) for 30 min. After the cells were washed with PBS, surface staining was performed using antibodies diluted in stain wash buffer (2% fetal bovine serum in PBS and [2 mM] EDTA) for 25 min on ice in the presence of 2.4G2 FcR blockade to reduce non-specific staining. For phenotype analysis cells were then fixed for 10 minutes using fixation/permeabilization solution (BD biosciences). For functional assays after fixable live/dead and surface staining, the cells were fixed and permeabilized for 1 h on ice in Fixation/Permeabilization solution (BD Bioscience), followed by intracellular staining in 1 X permeabilization wash buffer (BD Bioscience) with anti-IFNγ and anti-granzyme B (XMG1.2, NGBZ, eBioscience, Thermo Fisher Scientific) for 45 min. Antibodies used for surface staining were against: CD3 (17A2), CD49b (DX5), CD49a (HMα1), NKp46 (29A1.4), NK1.1(PK136), CD4 (RM4-5), CD8b (YTS156.7.7), KLRG1 (2F1/KLRG1), 2B4 (m2B4), Ly49I (YLI-90), Ly49H (3D10), CD94 (18d3), NKG2AB6 (16A11), LAG3 (C9B7W), PD1 (29F.1A12), PDL1(10F.9G2), CD107a (1D4B), CD45.1 (A20), CD45.2 (104). These antibodies were from Biolegend. Anti-Qa-1b (6A8.6F10.1A6) was from eBiosciences and anti-Ly49D (4E5) was from BD Biosciences. NKp46 (NCR1) ligand was stained using the soluble NKp46 receptor fused to human Fc (NCR1-hFc, RND systems). Bound soluble receptor was then detected using a secondary antibody anti-human IgG (). To assess apoptosis, cells were also stained using Annexin V (Annexin V staining kit, Biolegend). The cells were resuspended in 1 X PBS and analyzed using Guava easyCyte 12HT flow cytometer (Millipore-SIGMA) and FlowJo software (Tree Star).

### Cyst burdens

Cyst burdens were quantified using microscopy. Brains of mice infected with Type II strain ME49 were harvested and homogenized with a dounce homogenizer in 2 mls of 1 X PBS. 10 uls of homogenized brain was placed onto a microscope slide and covered with a cover slip. Microscope slides were examined and cysts in the homogenate were counted. A minimum of 5 slides per mouse was counted.

### Survival studies

WT B6 mice were infected with 10 cysts i.g. of the type II parasite strain ME49. After 5 weeks of infection mice were treated or not i.p. with 200 ug anti-NK1.1 (PK136, BioXCell) or 50 ug LEAF purified anti-NKp46 (Biolegend). Mouse treatments were performed every other day until completion of experiments. Mice were monitored daily for morbidity and mortality. Mice were evaluated on a 1-5 scale with 5 indicating highest morbidity. Mice reaching a level 5 score are not moving, severely hunched, eyes shut and not eating or drinking. Mice were sacrificed prior to death and after they reached level 5 clinical score for no more than 24 hours. For survival experiments after rechallenge, ME49 infected animals were treated or not with anit-NK1.1 (PK136, BioXcell). After the 2^nd^ dose of anti-NK1.1 mice were challenged with either a lethal dose 200 cysts of ME49 i.g. or 1000 tachyzoites of the type I highly virulent strain RH i.p. ILC depletion was continued every other day as in other experiments. Control mice were uninfected naïve B6 mice only given the challenge infection (either ME49 or RH). Survival of chronically infected rechallenged mice were monitored and assessed on the 1-5 scale as described above.

### Statistical analysis

Statistical analysis was performed using Prism 7.0d (GraphPad) and Microsoft Excel 2011. Significant differences were calculated using either unpaired Student’s t-test with Welch’s correction or analysis of variance (ANOVA). The log-rank (Mantel-Cox) test was used to evaluate survival rate. Data is presented in graphs as the mean± standard deviation (SD). Significance is denoted as follows: ns, not significant (p > 0.05) or significant with a maximum p-value of 0.05 or less.

## Results

### NK cell exhaustion

Previous studies have demonstrated that during late chronic *T. gondii* infection, CD4+ and CD8+ T cells develop immune exhaustion resulting in their dysfunction[10; 12; 13]. This ultimately results in the death of B6 mice in the late chronic stage of infection due to parasite reactivation. To further dissect the immune mechanisms contributing to T cell exhaustion during late chronic *T. gondii* infection, we investigated the role of innate lymphoid cells and more specifically NK cells. NK cells can participate in immune responses long after the innate response has transitioned into the adaptive response[20; 22]. NK cells acquire characteristics of memory. NK cells can also develop characteristics of immune exhaustion in the tumor microenvironment[27]. They traffic to tumor sites, have reduced numbers, effector function and upregulate PD1 expression on their surface. Based on the ability of NK cells to contribute to immunity after the innate response is over and their potential to develop immune exhaustion we determined whether NK cells were still in abundance during late chronic *T. gondii* infection and their immune exhaustion status. Mice were infected with 10 cysts of the Type II *T. gondii* strain ME49, known to induce T cell exhaustion during long-term infection. At week 5 and 7 post infection spleens were harvested and NK cell frequencies and numbers were measured using flow cytometry. Lineage negative (CD4-CD8-) cells were analyzed for CD49b+ cells. As shown in Figure 1A, the frequencies and absolute numbers of splenic NK cells (CD4-CD8-CD49b+) did not significantly decrease from week 5 to 7 post infection. As previously published week 7 is when CD4+ and CD8+ T cells decrease in both frequency and number[10; 13]. An increase in Programmed death 1 (PD1) on T cells is a hallmark of immune exhaustion. During late chronic *T. gondii* infection, both CD4+ and CD8+ T cells have been reported to increase their PD1 expression leading to loss of function of these T cells and parasite reactivation[10; 13]. To further assess whether NK cells exhibited characteristics of immune exhaustion during late chronic *T. gondii* infection we measured their PD1 expression. As shown in Figure 1B, the mean fluorescence intensity(MFI) of PD1 increased on both CD4+ and CD8+ T cells, however, NK cells did not increase their expression of PD1. In addition, the frequencies of CD4+ or CD8+ T cell PD1 high (PD1 Hi), PD1 intermediate (PD1 Int) both increased significantly at week 5 and 7 post infection (Figure 1C). The frequencies of PD1 Hi or Int did not change on NK cells over the course of infection. Another marker of exhaustion is lymphocyte activating gene 3 (LAG3) expression. LAG3 increases on CD4+ and CD8+ T cells during late chronic *T. gondii* infection[13]. We did not detect an increase in LAG3 expression (Figure 1 D)on NK cells during chronic *T. gondii* infection. Based on the results splenic NK cells do not appear to decrease in number or express PD1 or LAG3 at high levels compared to CD4+ and CD8+ T cells during chronic *T. gondii* infection.

**Figure 1.**
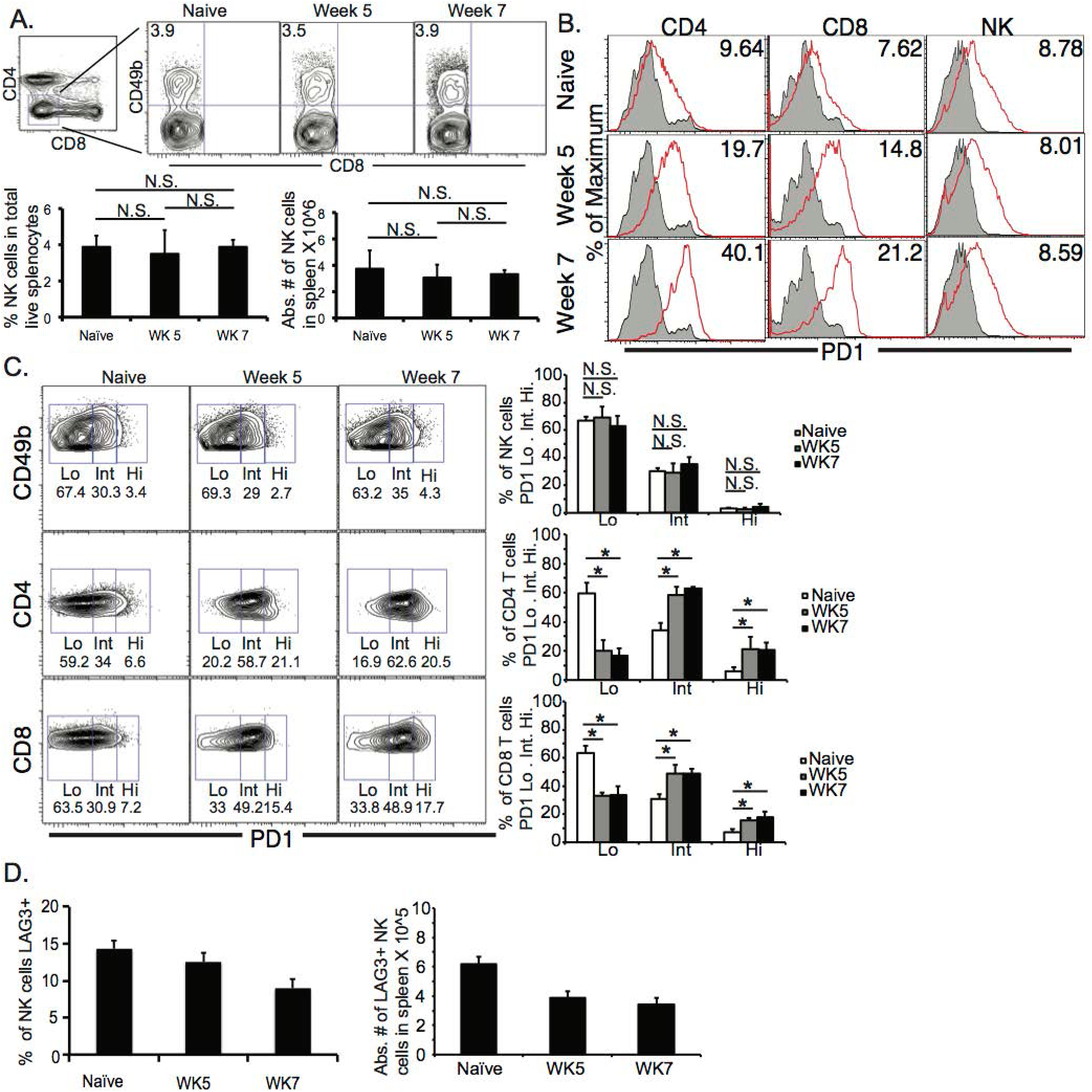
NK cells do not exhibit immune exhaustion characteristics during chronic *T. gondii* infection. C57BL/6 mice were orally infected or not with 10 cysts of ME49 and 5 and 7 weeks after infection spleen cells were analyzed for (A) NK cell (Lineage-CD49b+) frequency and number. NK cells, CD4+ and CD8+ T cells were then analyzed for (B) PD1 MFI and (C) PD1+ frequency based on separating the populations into PD1 low (Lo), intermediate (Int) and high (Hi). NK cells were then assayed for (D) LAG3 expression. Raw flow data presented are representative and based on means from 3 independent experiments. All graphs are mean ± SD. Graphs present pooled data from 2 independent experiments. Significance is denoted by * with a p≤0.05, n=4-5 mice per group.

### NK cell role in chronic *T. gondii* infection

Based on results of our studies, NK cells did not appear to develop some characteristics of immune exhaustion raising the question about how might NK cells contribute to immune control of *T. gondii* during chronic infection. WT B6 mice typically succumb to spontaneous reactivation of the parasite in the CNS and die[10]. Cyst reactivation in the brain can be observed via parasitemia and a decrease in cyst number in the brains of late chronic infected mice. Interestingly, blockade with anti-PDL1 antibody appears to rescue these animals from death and slow down parasite reactivation resulting in maintenance of cysts in the CNS[10; 11; 12]. To begin to address how NK cells are behaving during chronic *T. gondii* infection NK cells were depleted using anti-NK1.1 in mice starting at week 5 post infection. Mice were treated every other day until the experiment was terminated at 100 days post infection. Mice with NK cells began to succumb to the infection around week 5 (49 days) post infection (Figure 2A). Mice treated with anti-NK1.1 did not start to succumb to the infection until 80 days post infection. All mice with NK cells were dead by 80 days post infection whereas 50 % of mice depleted of their NK cells were still alive at 100 days post infection (Figure 2A). We next measured whether cyst burdens in the brain were maintained better when NK cells were depleted. As shown in Figure 2B, mouse brain cyst burdens were higher in mice that were depleted of NK cells than mice with NK cells. To fully test whether NK cells were inhibiting immune control of the parasite, B6 mice were infected with ME49 and at 5 weeks depleted of their NK cells or not with anti-NK1.1. 2 days after start of treatment, mice were challenged with a lethal dose of either 200 cysts of ME49 i.g. or 1000 tachyzoites of RH i.p. and monitored for survival. As shown in Figure 2 C, mice with their NK cells succumbed to challenge significantly earlier than mice with NK cells depleted. This result demonstrates that NK cells appear to have a negative affect on long-term immunity to chronic *T. gondii* infection by promoting parasite reactivation and reducing the effectiveness of adaptive recall responses.

**Figure 2.**
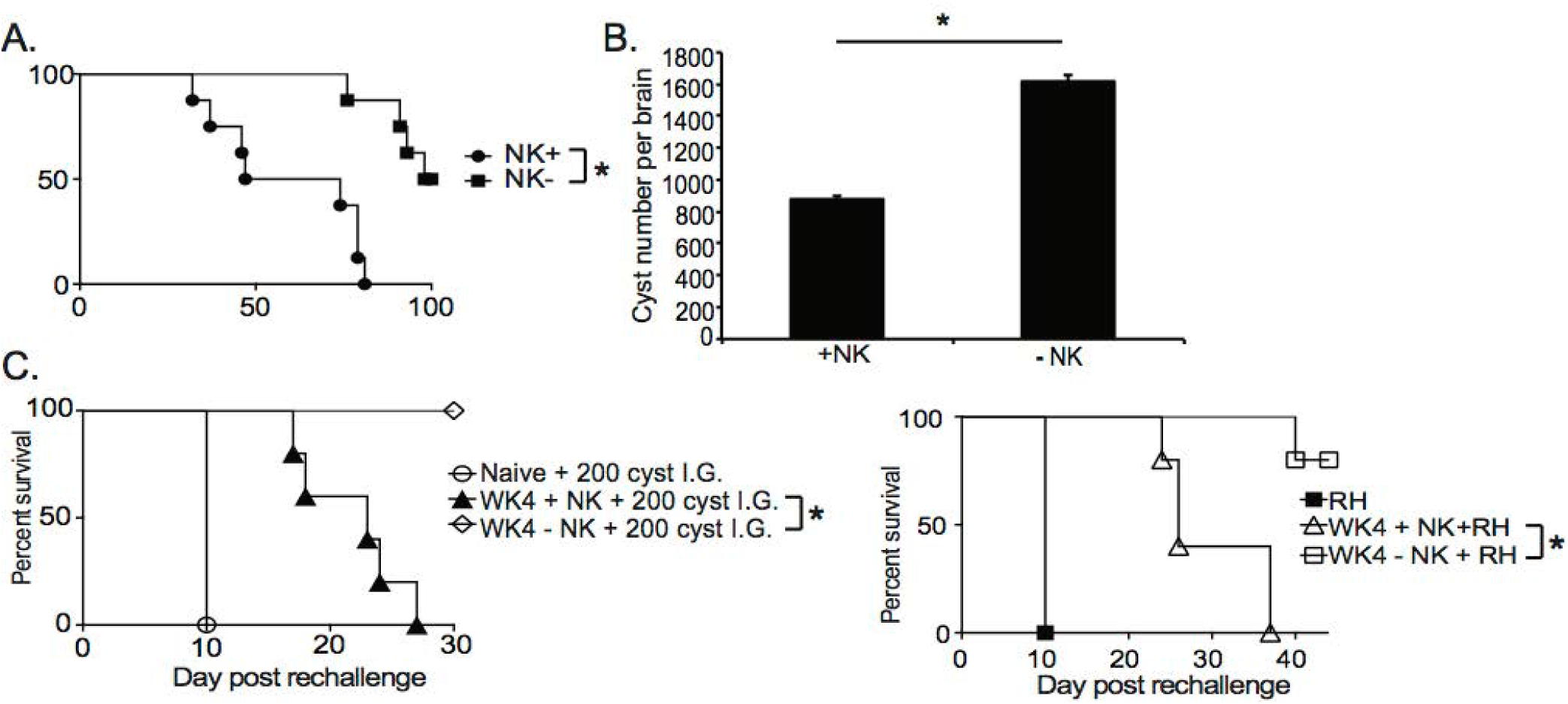
NK cells promote immune exhaustion and parasite reactivation during chronic *T. gondii* infection. C57BL/6 mice were infected orally with 10 cysts of ME49 and infection outcome monitored. (A) Survival after 200 ug anti-NK1.1 every 2^nd^ day starting at week 5 p.i. (B) Brain cyst burden after anti-NK1.1 treatment starting at week 5 p.i. (C) Survival of chronically infected animals after lethal secondary 200 ME49 cyst or 1000 tachyzoite RH strain challenge with or without anti-NK1.1. The log-rank (Mantel-Cox) test was used to evaluate survival rates. Ordinary one-way ANOVA was used to evaluate the parasite burdens. All graphs are mean ± SD. The data presented in graphs is pooled from 3 independent experiments with an n=4-5 per group. Total number of mice used was 12-15 animals per group. * denotes significance as p≤ 0.05.

### NK cell impact on CD8+ T cell responses

Parasite reactivation and mouse death during chronic *T. gondii* infection occurs when parasite specific CD8+ T cells develop immune exhaustion[10]. Their exhaustion results in decreased frequency and number of polyfunctional (IFNγ+CD107a+, or IFNγ Granzyme B+) CD8+ T cells in the periphery and CNS. Since depletion of NK cells during chronic *T. gondii* resulted better survival we measured how this impacted polyfunctional CD8+ T cells in the spleen and brain. Infected mice were treated starting at week 5 post infection with anti-NK1.1 for 2 weeks as in previous experiments and spleen and brain cells were assayed for the frequency and absolute number of polyfunctional CD8+ T cells. As shown in figure 3A, NK cell depletion of week 5 infected mice resulted in the maintenance of the frequency and absolute number of IFNγ+CD107a+ CD8+ T cells. Similarly, in the brain, NK cell depletion starting at week 5 post infection significantly increased the frequency of IFNγ+GrzB+ CD8+ T cells. Several studies have demonstrated that during acute viral infections (MCMV, LCMV) infection, NK cells are negative regulators of priming of adaptive immune responses[29; 30; 32; 33; 37; 38; 39]. This negative regulation promotes viral persistence and immune exhaustion of the T cells. However, during acute *T. gondii* infection previous studies suggest that NK cells could be positive regulators of the priming of adaptive immune responses against the parasite[40; 41; 42; 43]. Many of the earlier *T. gondii* studies used anti-asialo GM1 antibody to deplete NK cells without knowing that this antibody targets not only NK cells, but effector populations of CD8+ T cells [44]. We tested whether NK cells provided a different function during acute *T. gondii* infection and promoted priming of CD8+ T cells as compared to NK cells in chronic *T. gondii* infection, which appear to inhibit CD8+ T cell function. B6 mice were treated or not with anti-NK1.1 1 day prior to infection with ME49 strain of *T. gondii* and then infected with 10 cysts i.g. NK depleted mice were treated with anit-NK1.1 for 6 days and on day 7 all mice were harvested and their spleen CD8+ T cell functionality was measured. As shown in figure 3D and E, CD8+ T cells were activated by day 7 post infection, however, NK cell depleted animals had significantly fewer activated CD8+ T cells (IFNγ+CD107a+ plus IFNγ+CD107a-) than mice that still have their NK cells. Thus, during acute infection NK cells are important for priming CD8+ T cells to protect against infection, but during chronic *T. gondii* infection, NK cells change their function and negatively regulate CD8+ T cells to promote parasite reactivation and mouse death. To further define why polyfunctional CD8+ T cells were reduced during chronic *T. gondii* infection in the presence of NK cells, we measured CD8+ T cell apoptosis using Annexin V staining in the presence or absence of NK cells. As presented in figure 4A, apoptosis was significantly increased in chronically infected mice at week 5 and 7 post infection. NK cell depletion significantly reduced the level of CD8+ T cell apoptosis in chronically infected mice at week 7 post infection. Similar results were obtained by measuring active caspase 3/7 in CD8+T cells (data not shown). NK cells appear to contribute to parasite reactivation and CD8+ T cell exhaustion during chronic *T. gondii* infection by increasing CD8+ T cell apoptosis.

**Figure 3.**
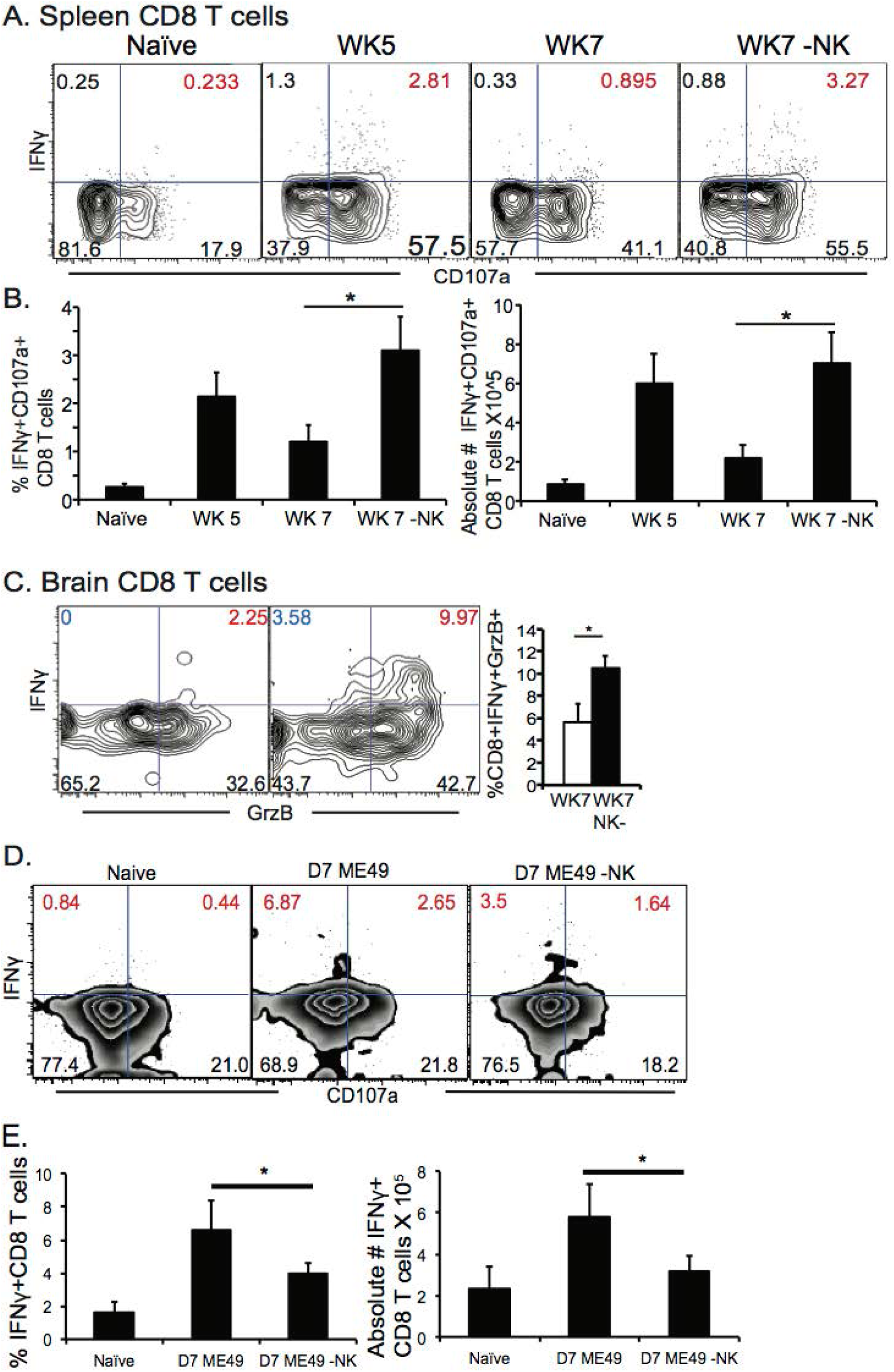
NK cells reduce polyfunctional CD8+ T cells by increasing their apoptosis during chronic *T. gondii* infection. C57BL/6 mice were infected as described and 5 weeks after infection treated with anti-NK1.1 i.p. Brain and spleen cells were harvested and restimulated *ex vivo* with TLA. (A) Contour plots present spleen CD8+ T cells analyzed for IFNγ+ X CD107a+. (B) Graphs present frequency and absolute # of polyfunctional IFNγ+CD107a+CD8+T cells. (C) Contour plots of brain IFNγ+GrzB+CD8+ T cells (red numbers are frequency) are presented. (D and E) B6 mice were treated or not with 200 ug of anti-NK1.1 starting at D-1 then infected with 10 cysts of ME49 strain i.g. Mice were treated every other day with anit-NK1.1. On day 7 post infection animals were sacrificed and spleen cells isolated then stimulated with TLA and stained for polyfunctionality (IFNγ+CD107a+). (D) Contour plots present CD8+ T cells stained for IFNγ+ X CD107a+ during acute *T. gondii* infection. (E) Graphs present pooled data from 2 experiments showing frequency (%) and absolute number of IFNγ+ CD8+ T cells in spleen. All graphs are mean ± SD. Experiments were repeated 3-5 independent times with an n=4 mice per group. * denotes significance with a p≤0.05.

**Figure 4.**
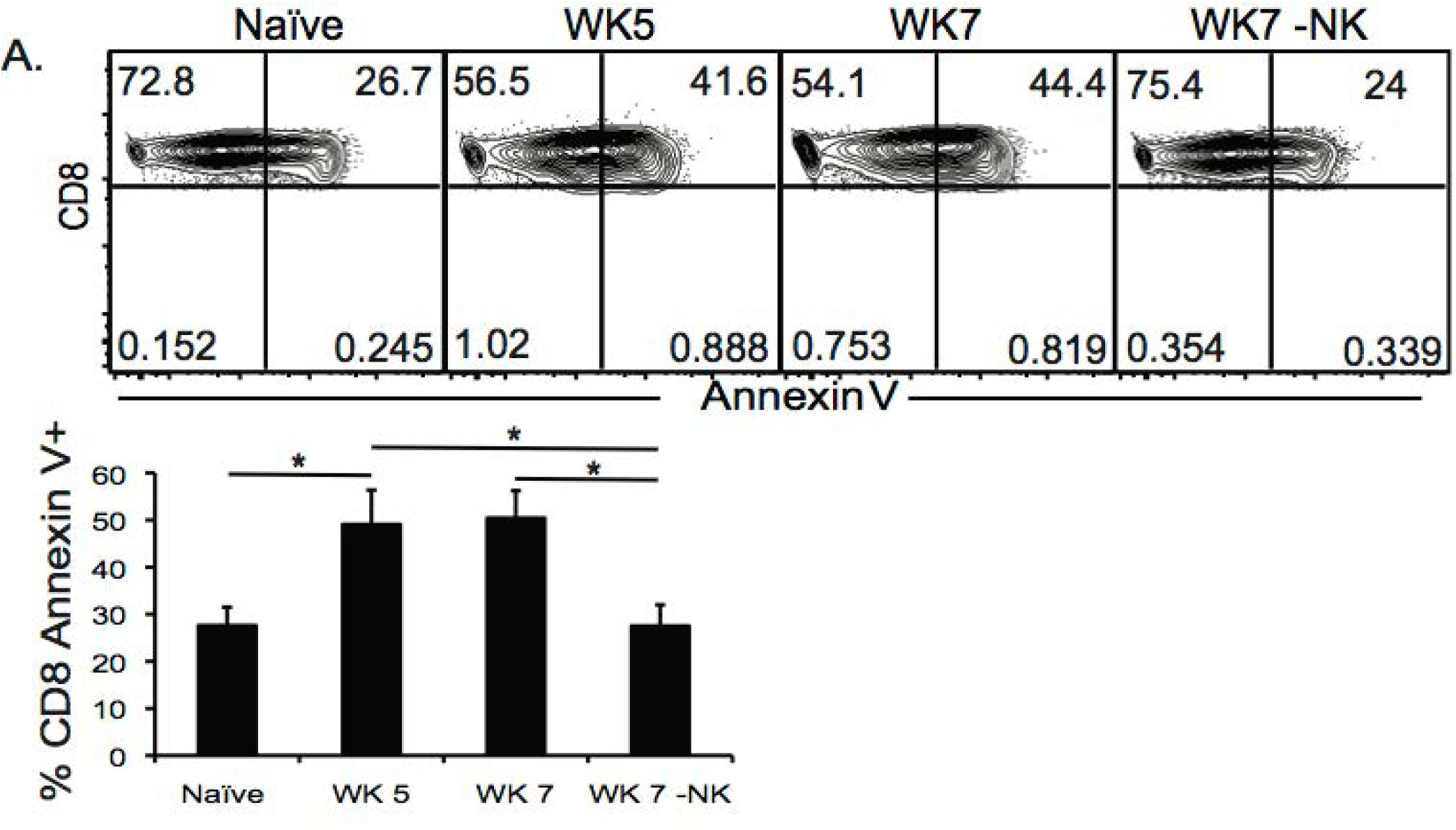
NK cells increase CD8+ T cells apoptosis during chronic *T. gondii* infection. C57BL/6 mice were infected and some groups at week 5 after infection were treated or not with anti-NK1.1 i.p. as described. Week 5, week 7 non treated and week 7 treated infected mice were sacrificed and spleen cells isolated for Annexin V staining and compared to naïve animals. (A) Contour plots present frequency of Annexin V+ CD8+ T cells and graphs present mean ± SD. Data presented are from 1 experiment repeated 3 independent times. Significance is denoted by * with a p≤0.05, n=4-5 mice per group.

### NK cell phenotype during chronic T. gondii infection

Published studies indicate that NK cells use several mechanisms to regulate adaptive immune responses[29; 31; 32; 33; 34; 45; 46; 47]. Many of these mechanisms rely on the expression of specific NK cell receptors, which allow the NK cells to target specific adaptive immune cell populations and induce their apoptosis, lysis or suppression. NK cell receptors are also very important for normal protective NK cell functions including Ly49H, which recognize m157 of MCMV[22; 48]. These specific interactions promote the enrichment of NK cell subpopulations expressing specific receptor combinations as was demonstrated for memory-like NK cells during MCMV infection. During acute *T. gondii* infection, we have previously published that there does not appear to be a dominant NK cell population activated. *T. gondii* infection may only induce cytokine dependent NK cell activation resulting in global activation of a large array of different IFNγ producing and protective NK cell subpopulations [49]. In the studies presented here, NK cells appear to change their function to promote CD8+ T cell compartment dysfunction and parasite reactivation. This suggests the NK cell compartment could be modified and a specific NK cell subpopulation develops to erode immunity to the parasite during chronic *T. gondii* infection. To begin to define what NK cell receptors might be involved in contributing to CD8+ T cell exhaustion during chronic *T. gondii* infection, we performed an exhaustive assessment of NK cell receptor expression during week 5 chronic *T. gondii* infection. As shown in figure 5A, the NK cell compartment had significantly reduced frequencies of cells that expressed 2B4, Ly49H, Ly49D and Ly49I. This was observed on lineage – CD49b+ cells in the spleen. We did not detect any differences in TRAIL expression (data not shown). We observed significant increases in the frequencies of NK cells (lin-CD49b+) that expressed KLRG1 and NKG2A (Figure 5A). The number of NK cells expressing KLRG1 also increased significantly during chronic *T. gondii* infection. The significant increase in KLRG1+ NK cells and NKG2A+ NK cells suggested these cells were being enriched within the NK cell compartment. A recent study suggests that CD49a+ ILC1 may develop from NK cells and exhibit this phenotype in the liver of mice infected with the vaccine strain of *T. gondii cps1-1* and a different limited cyst forming type II strain Prugniaud [50]. Therefore to determine if this was also occurring in NK cells in the spleens after infection with the vaccine strain, we infected mice with *cps1-1* and 5 weeks later assayed the spleen cells for CD94+NKG2A+ NK cells. As shown in figure 5B, *cps1-1* strain parasites did not induce the same increase in frequency of CD94+ NKG2A+ NK cells as did ME49. To further investigate whether ILC1 were enriched in the NK cell compartment, we measured the frequencies of ILC1 (CD49a+CD49b-) compared to NK cells (CD49a-CD49b+) in the CD3- and CD3-NKp46+ cell populations in the spleens of chronically infected mice. As shown in figure 5C, NK cells (CD49b+CD49a-) comprised 22% of the CD3-population of cells and 60% of the lineage negative NKp46+ population of cells. ILC1 were present at a very low level. Therefore in the spleen, NK cells appear to be the dominant population of ILC present and they are enriched for a specific receptor phenotype, which is Lin – CD49b+ CD49a-NKp46+ CD94+ NKG2A+KLRG1+.

**Figure 5.**
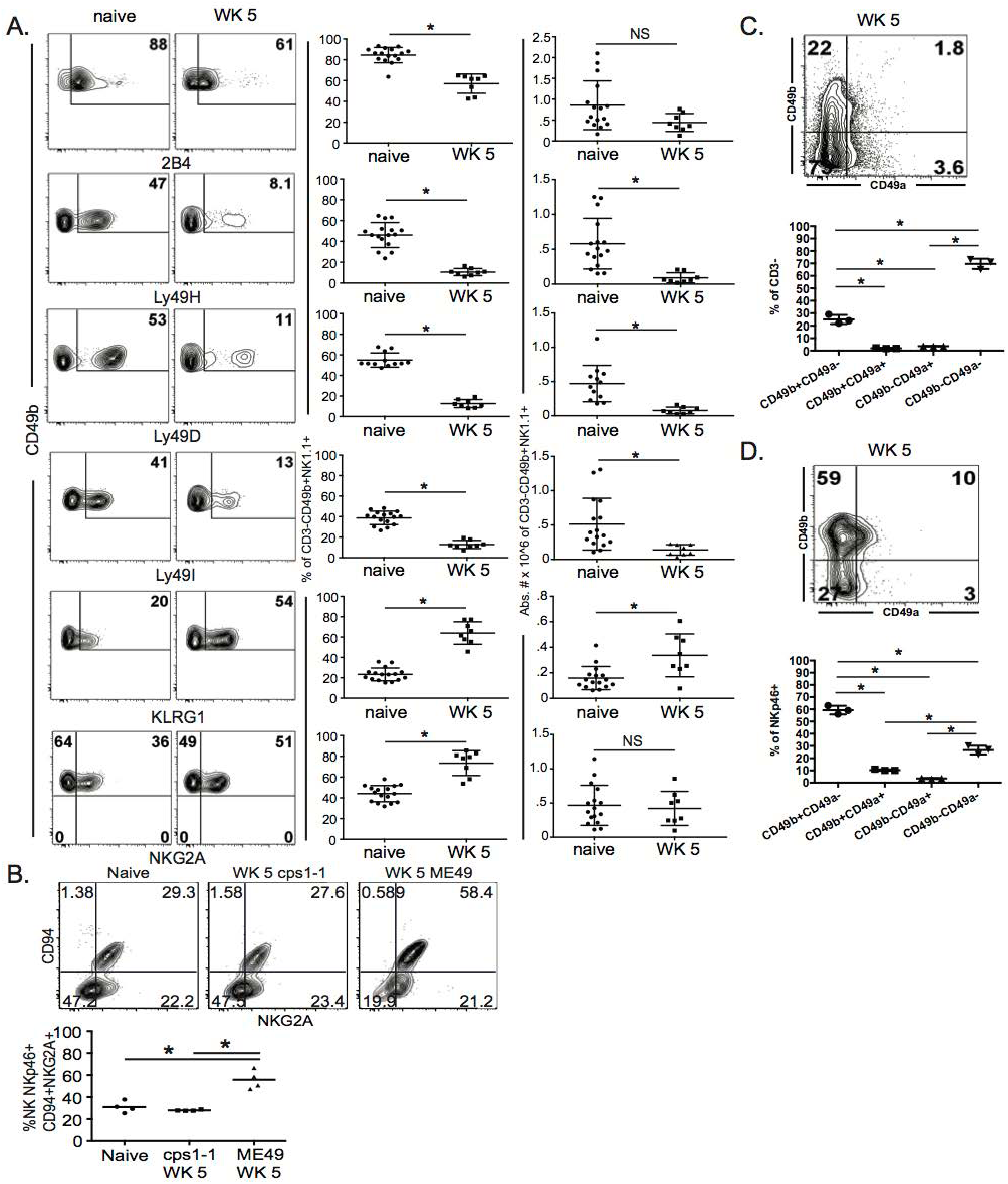
NKp46+NKG2A+KLRG1+ NK cells are enriched in spleen during chronic *T. gondii* infection. C57BL/6 mice were orally infected with 10 cysts of ME49 and analyzed for the NK cell receptors 2B4, Ly49H, Ly49D, Ly49I, KLRG1, CD94 and NKG2A by flow cytometry. (A) Contour plots present the frequency of CD49b+ X receptor + cells comparing naïve animals to week 5 post infection. Graphs show pooled data from 3 experiments of frequency and absolute number of CD49b+ Receptor+ cells. (B) Mice were infected with either 1 × 10^6^ tachyzoites of *csp1-1* i.p. or 10 cysts ME49 i.g. At week 5 post infection, lineage-CD49b+NKp46+ cells were analyzed for CD94 X NKG2A. Contour plots present data from one experiment showing frequency of CD94 X NKG2A cell populations. The graph presents data from 1 experiment comparing the frequency of CD94+NKG2A+ cells between naïve, *cps1-1* and ME49 mice. (C) Mice were infected with 10 cysts of ME49 i.p. then spleen cells were analyzed at week 5 post infection for CD49a X CD49b to identify ILC1 compared to NK cells. Contour plot presents the frequency of CD49a X CD49b cells in the CD3-population. (D) Contour plot presents the frequency of CD49a X CD49b cells in the CD3-NKp46+ population. (C-D) Graphs present the frequency of CD49b+CD49a-, CD49b+CD49a+, CD49b-CD49a+ and CD49b-CD49a-. Experiments were repeated independently a minimum of 2 times with n=3-4 per group. * denotes significance with a p≤0.05.

### NK cell function during chronic T. gondii infection

NK cells are the cytotoxic cells of the ILC lineage[51; 52]. NK cells are also capable of producing high levels of IFNγ upon activation. ILC1 are not cytotoxic and produce high levels of IFNγ upon activation. During acute *T. gondii* infection NK cells and ILC1 are known to produce IFNγ in an IL-12 dependent manner[16; 18; 19; 53; 54] We have recently demonstrated that after vaccination, NK cells respond a second time to help control challenge infection by producing IFNγ in an IL-12 and IL-23 dependent manner [55]. We observe that NK cells during chronic *T. gondii* infection modify their role in immunity to the parasite and are not protective, but detrimental. They also express an altered receptor repertoire that suggests enrichment for a specific cell phenotype. Therefore to begin to investigate how NK cells are negative regulators of CD8+ T cells during chronic *T. gondii* infection we first assayed their function. Mice were infected as above and starting at week 5 post infection we assessed NK cell (CD3-CD49b+ NKp46+) function (IFNγ X CD107a) by flow cytometry. As shown in figure 6A, after *ex vivo* stimulation, naïve NK cells were capable of producing both IFNγ and expressing the surrogate cytotoxicity marker CD107a. However, after week 5 of infection (Figure 6a), NK cells produced very little IFNγ while significantly increasing their CD107a expression. This pattern of function was observed also at week 7 post infection. Increases in CD107a+ NK cells were observed in both frequency and absolute number. Interestingly, the frequency of IFNγ+ NK cells continued to decrease from week 5 to week 7 post infection. The data shown in figure 6A was generated *ex vivo* by using plate bound anti-NK1.1 crosslinking. We repeated *ex vivo* analysis using PMA/Ionomycin and still the NK cells did not produce IFNγ (data not shown). Interestingly, if these cells were ILC1, we would have expected them to produce IFNγ. We next measured whether the NK cells expressed PD-L1, the ligand for PD1. As shown in Figure 6B, splenic NK cells did not appear to increase their expression of PD-L1 as PD-L1 MFI was not significantly different between week 5 and 7 post infection. During acute systemic *T. gondii* infections, NK cells have been shown to produce IL-10[34]. Therefore, we obtained IL-10GFP TIGER reporter mice and infected them with 10 cysts of ME49 i.g. Comparing naïve to week 5 post infected mice (Figure 6C), we did not observe any IL-10 production by NK cells. A recent study demonstrated that NKp46+ ILC could contribute to the development of neurodegenerative disease by being in the CNS and promoting Th17 responses[47]. We next determined whether there were NK cells in the CNS. Mice were infected as previously described and at week 5 post infection, mice were perfused, brains dissected and immune cells isolated. Cells were analyzed for CD3-NKp46+ populations. As shown in figure 6D, we did not observe an increase in frequency of CD3-Nkp46+ NK cells in the CNS of *T. gondii* chronically infected mice at week 5 post infection. Our investigation of the function of the NK cells acting as negative regulators of immunity during chronic *T. gondii* infection suggests that NK cells have reduced IFNγ production, but may increase cytotoxicity. They do not produce IL-10 and they are functioning from outside the CNS to cause CD8+ T cell dysfunction.

**Figure 6.**
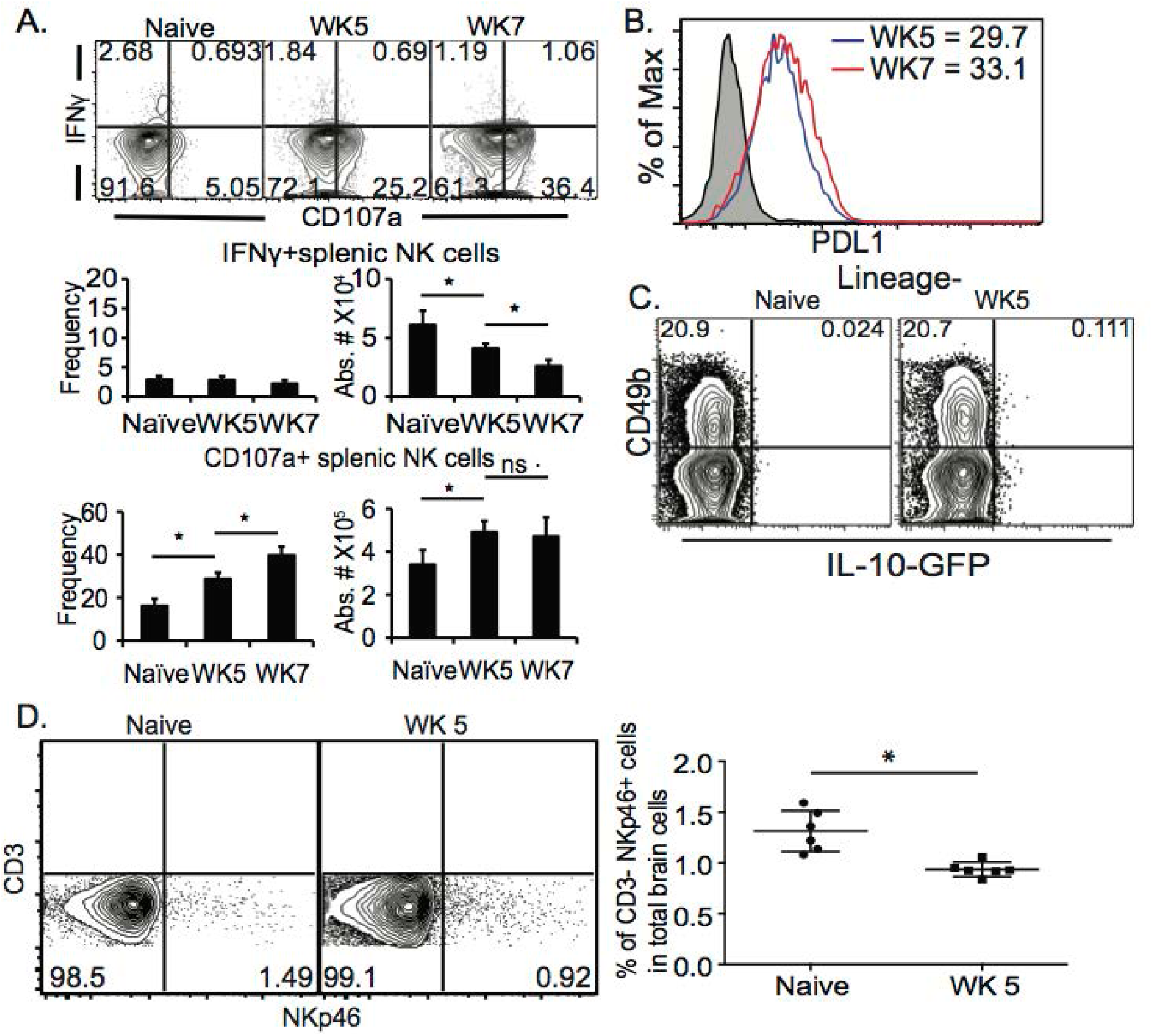
NK cells have altered function during chronic *T. gondii* infection. C57BL/6 or IL-10 reporter TIGER-GFP mice were orally infected with 10 cysts of ME49 and 5 and/or 7 weeks after infection spleen cells analyzed for function. (A) Spleen cells were stimulated *ex vivo* with plate bound anti-NK1.1 then stained for NK cells (CD3-CD49b+ NKp46+) IFNγ and CD107a. Contour plots present frequency data gated on NK cells and compares IFNγ X CD107a. Graphs present the frequency and absolute number of IFNγ+ NK cells (top graphs) and CD107a+ NK cells (bottom graphs). Graphs present mean ± SD. (B) Splenic NK cells were assayed for PDL1 expression. Histogram presents the MFI of PD-L1 on NK cells from week 5 and 7 post infection mice. (C) Contour plots present the frequency of IL-10 GFP+ NK cells in naïve compared to week 5 post infection mice. (D) Brain cells were isolated and stained for lineage markers, CD49b and NKp46. Contour plots present the frequency of CD3-NKp46+ cells in the CNS. Graphs present the pooled data from 2 experiments of frequency of CD3-NKp46+ cells in the CNS. Experiments were repeated at least 2 independent times with an n=3-5 mice per group. Significance is denoted by * with a p≤0.05.

### NKp46 and NKG2A NK cells during chronic *T. gondii* infection

The NK cell phenotype we observed Lin – CD49b+ CD49a-NKp46+ CD94+ NKG2A+KLRG1+ suggest that NKp46 and NKG2A may contribute NK cell negative regulation of the immune response to *T. gondii* in chronically infected mice. This is based on the concept of NK cell licensing[56]. NK cell licensing determines the responsiveness of NK cells to self versus non-self. A licensed NK cell expresses both activating and inhibitory receptors on its surface and as a result is tuned or permitted to respond when self is absent. An absence or reduction in self, usually reduced MHC expression can be detected on a target cell. At the same time, increases in non-self detected by elevated ligands binding to the activating receptor activate the NK cell. NKp46 is an activating receptor expressed on NK cells, ILC1 and some ILC3[52; 57]. The ligand NKp46 recognizes is not very well described. Potential ligands for NKp46 vary in source and structure and to date may include Influenza virus HA, Sigma 1 protein of Reovirus and *Candida glabrata* proteins Epa 1, 6 and 7[58; 59; 60]. NKp46 is a natural cytoxicity receptor, also called NCR1 and is known once it engages its ligand (NCR1-ligand) to lyse target cells[35]. NKp46 can also promote the expansion and survival of NK cells similar to other activating receptors[35; 61]. NKG2A is an inhibitory receptor that recognizes non-classical MHC Class I known as Qa-1b[62; 63]. NKG2A prevents NK cell activation. Based on the licensing paradigm and our data we hypothesized that during chronic *T. gondii* infection, there was an increase in non-self (NCR1-ligand) while there was a decrease in self (Qa-1b) which in turn caused NK cells to negatively regulate CD8+ T cells resulting in parasite. reactivation and death. To test this hypothesis spleens from chronically infected mice at week 5 and 7 post infection were isolated and the expression levels of NKp46-ligand and Qa-1b were measured on total splenocytes and CD8+ T cells and compared to naïve animals. NCR1-ligand was detected using soluble murine NCR1 (NKp46) fused to human Fc and Qa-1b using anti-Qa-1b antibody. In naïve mice total splenocytes were positive for Qa-1b and largely negative for ligands that were bound by NCR1(Figure 7A, top and bottom panels, respectively). At week 5 post infection Qa-1b was significantly increased in expression and NCR1-ligand remained low compared to naïve mice. At week 7 post infection Qa-1b expression was decreased significantly compared to week 5 and naïve animals while NCR1-ligand was increased significantly (Figure 7A). As shown in figure 7B, this pattern of Qa-1b and NCR1-ligand expression was similar when CD8+ T cells were gated and assessed. However, the changes in Qa-1b and NCR1-ligand did not appear to be greater on CD8+ T cells than total splenocytes. We performed preliminary assessments of whether the NK cells were actually more cytotoxic, but did not find any significant increase (data not shown), suggesting these interactions were promoting NK cell survival and maturation as measure by KLRG1 expression on the NK cells (Figure 5A). Overall the decrease in self (Qa-1b) and the increase in non-self (NCR1-ligand) support the concept that NK cell licensing was contributing to CD8+ T cell dysfunction in some way. Therefore to test that these NK cells via a licensing process were contributing to immune dysfunction during chronic *T. gondii* infection, we infected animals as before and starting at week 5 post infection we treated or not mice with non-depleting anti-NKp46 blocking antibody[35]. This approach would not deplete NK cells, but simply block the interaction between NKp46 and the unknown ligand thus potentially decrease NK cell negative regulation of CD8+ T cells. As shown in figure 7C, anti-NKp46 significantly prolonged the life of mice with chronic *T. gondii* infection compared to no treatment controls. These results suggest that modifications of self versus non-self and NK cell recognition of these modifications via NKp46 and NKG2A receptors potentiate NK cell dependent negative regulation of CD8+ T cells responses during chronic *T. gondii* infection. NK cells as a result contribute to immune exhaustion not early during infection, but later after chronic *T. gondii* infection is established.

**Figure 7.**
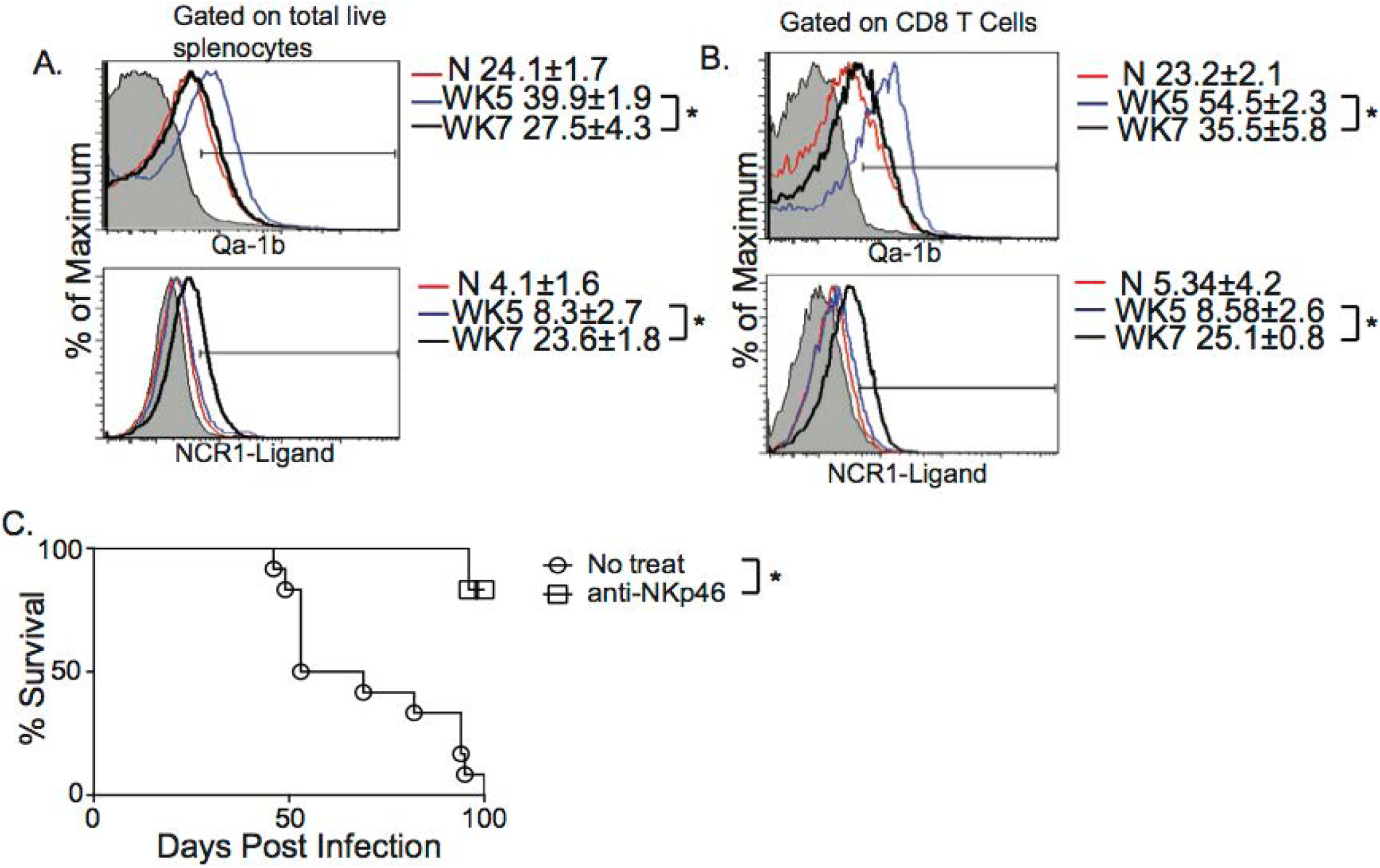
Blockade of NKp46 rescues mice from death caused by CD8+ T cell exhaustion induced parasite reactivation. C57BL/6 mice were orally infected as above and total splenocytes were assayed for NKG2A ligand Qa-1b and NKp46 ligand using soluble NCR1 fused to human Ig Fc. (A) Histograms present the MFI ± SD of QA-1b (top) and NCR1-ligand (bottom) from total splenocytes. (B) Histograms present the MFI ± SD of QA-1b (top) and NCR1-ligand (bottom) from CD8+ T cells. (C) Mice were infected with 10 cysts of ME49 i.g. and treated or not with 50 ug of anti-NKp46 i.p. starting at week 5 p.i. Mice were treated every other day for the duration of this experiment. The survival graph presents pooled data from 3 independent experiments. All experiments were repeated a minimum of 2 times. The log-rank (Mantel-Cox) test was used to evaluate survival rates. * denotes significance with p≤0.05.

## Discussion

The immune mechanisms regulating CD8+ T cell exhaustion resulting in reactivation of chronic *T. gondii* infections are poorly understood. In this study we sought to further explore these mechansisms and proposed that NK cells could contribute to this process. NK cells are innate immune cells and belong to a growing family of immune cells known as innate lymphoid cells[52; 64]. NK cells provide a first of defense against many pathogens via their ability to lyse tumor cells and infected cells and produce high levels of IFNγ. Although they have a primary role in innate immune protection, they can also contribute to long-term immunity. NK cells participate in memory responses by further differentiating and developing long life and more efficient recall responses[20; 21; 22]. During acute viral infections, systemic infections and in the tumor microenvironment NK cells can dysregulate CD4+ and CD8+ T cell responses promoting pathogen and tumor persistence and immune exhaustion [28; 29; 30; 32; 33; 34; 37; 38; 46]. In addition NK cells can become exhausted themselves in different tumors models and infection[26]. Based on this published knowledge of the complexity of NK cell biology, we tested whether NK cells become exhausted during chronic *T. gondii* infection, how they impact long term immunity to the chronic stage of infection and the mechanisms involved. Our studies demonstrate that NK cells do not appear to become exhausted because their numbers are stable and they do not increase PD1 or LAG3 expression despite losing the ability to produce IFNγ. They appear to enhance parasite reactivation and erode secondary immune responses in chronically infected animals. They accomplish this by reducing CD8+ T cell function by increasing their apoptosis. NK cells have increased activation as indicated by high KLRG1 and CD107a expression. During chronic *T. gondii* infection NK cells develop a unique Lin-CD49b+CD49a-Ly49-NKp46+CD94+NKG2A+ phenotype suggesting that these cells receive signals from altered self through NKp46 recognition of specific ligands and a reduction in Qa-1b. Indeed staining of total spleen and CD8+ T cells with soluble NCR1 and anti-Qa-1b indicate there is a significant change in altered self during chronic *T. gondii* infection. Our studies further support this hypothesis when we block NKp46 interaction and rescue chronically infected mice from death caused by CD8+ T cell exhaustion and parasite reactivation similarly to depletion of NK cells. Overall we find that NK cells are essential for acute immune protection by helping to control the parasite with IFNγ and also by helping to prime CD8+ T cells. However, during chronic *T. gondii* infection NK cells develop a response that contributes to CD8+ T cell dysfunction thereby promoting parasite reactivation in mice. NK cells can develop immune exhaustion in the tumor microenvironment, after overstimulation and during HCV infection[24; 25; 26; 27]. Our results suggest that NK cells are not becoming exhausted, but are developing into cells that negatively regulate the CD8+ T cell responses during chronic *T. gondii* infection. CD8+ T cells are known to develop immune exhaustion during chronic *T. gondii* infection[10; 12]. This leads to the reactivation of encysted parasites in the CNS and ultimately results in death of B6 mice. CD8+ T cell exhaustion during chronic *T. gondii* infection is marked by reduced CD8+ T cell numbers, decreased frequencies and numbers of IFNγ+CD8+ T cells in the spleen and brain, increased CD8+ T cell apoptosis and high expression of PD1 on the surface of CD8+ T cells. These are hallmarks of CD8+ T cell exhaustion in several infection and disease models[14]. Our results demonstrate that NK cells are present in the spleen during chronic *T. gondii* infection, they do not have reduced numbers and do not express high levels of PD1 or LAG3 as compared to CD4+ and CD8+ T cells. When we investigated NK cell function (IFNγ and CD107a), we observed that although NK cells in chronic *T. gondii* infection lose the ability to produce IFNγ, they increase their CD107a expression indicating a gain of function. Moreover, NK cell depletion rescued mice from death by helping restore CD8+ T cell function in spleen and brain, helped maintain encystation of the parasite and enhanced the survival of chronically infected mice after secondary parasite challenge. Although, NK cells may lose the ability to produce IFNγ, our results suggest that unlike tumor, overstimulation and persistent HCV infection[24; 25; 26; 27], they are also gaining function that negatively regulates the adaptive response to chronic *T. gondii* infection. The mechanism by which they are causing this negative regulation is unclear and will be important in future studies.

NK cells in the steady state express a stochastic array of activating and inhibitory receptors that help regulate their function[65; 66; 67]. In mice this includes the Ly49 family of receptors (D-I), natural cytotoxicity receptors (NCRs), NKG2D, 2B4 and CD94/NKG2A. In the naïve state in B6 mice, these receptors are expressed on most NK cells in different combinations, but at relatively high frequencies. Our data demonstrates that NK cells during chronic *T. gondii* infection have altered expression of NK cell receptors. We observe a near complete loss of Ly49 D, H and I. At the same time we observed the maintenance of CD49b+ NKp46+ NK1.1+ cells. Within this population the frequency of CD94+NKG2A+ cells increased dramatically and this increase was only observed during persistent chronic *T. gondii* infection and not after infection with the non-persistent vaccine strain. These cells appear to have a licensed NK cell phenotype because of the presence of both activating and inhibitory receptors on their surface[56]. Thus the licensing paradigm could explain why this phenotype of NK cells develops during chronic *T. gondii* infection. In this situation, the chronic infection environment in Toxoplasmosis causes the activating receptor NKp46 to recognize a ligand expressed on target cells that potentiates the activation, survival and increased abundance of NKp46+ CD94+NKG2A+ NK cells. This is what occurs with other activating receptors including Ly49H after it recognizes m157 from MCMV[22; 48]. As a result of Ly49H and m157 interaction, Ly49H+ NK cells are more abundant, have longer life and can respond more efficiently to secondary infection. While Ly49H interaction with MCMV m157 could directly activate all Ly49H positive NK cells regardless of inhibitory receptor expression, our data suggests that because of the higher frequency of CD94+NKG2A+ NK cells within the NKp46+ population, that loss of the inhibitory signal through NKG2A also helps promote the development of this NK cell population. NKG2A recognizes the non-classical MHC protein Qa-1b[63]. We observe that Qa-1b is decreased in expression by week 7 post parasite infection. This could take the brakes off of the NK cells and upon interaction of NKp46 with NCR1-ligand results in the enrichment of this phenotype of NK cells and their activation during chronic *T. gondii* infection. What NKp46 could be recognizing is still a mystery during chronic *T. gondii* infection. The ligands for NKp46 vary in source and structure and to date may include Influenza virus HA, Sigma 1 protein of Reovirus and *Candida glabrata* proteins Epa 1, 6 and 7[58; 59; 60]. Our data indicate that there is increased staining of spleen cells and CD8+ T cells with soluble NCR1. What protein modifications are occurring or genes that are being expressed to produce this ligand are unclear, however, the increase in binding of soluble NKp46 supports the hypothesis that the NKp46 signal is required for the development of this unique NK cell population during chronic *T. gondii* infection.

Another important phenotype we observe is the increase in KLRG1+ NK cells in chronically infected mice. KLRG1 is an inhibitory receptor expressed more highly as NK cells mature[15; 68]. NK cell maturation is activating receptor dependent. Recent studies investigating exhausted NK cells during chronic stimulation suggest that increased KLRG1 indicates NK cell exhaustion [25]. In this study, NKG2D interaction with high levels of NKG2D ligands results in increased KLRG1 expression and loss of NKG2D expression on the cells and NK cell exhaustion. Therefore another possible explanation of the development the phenotype of NK cells during chronic *T. gondii* infection could be that ligands for other receptors are highly upregulated during chronic *T. gondii* infection. This could then explain why we observe a loss of expression of Ly49 D, Ly49H, and Ly49I+ NK cells. However, we performed an exhaustive analysis of known murine NK cell receptor ligands and we did not detect any increase in their expression during chronic *T. gondii* infection (data not shown). We only observed increases in NKp46-ligand and reduced Qa-1b expression. Moreover, blockade of NKp46 with a non-depleting anti-NKp46 antibody rescued mice to a similar level from death compared to NK cell depletion with anti-NK1.1. A recent study demonstrates that NK cells are plastic during *T. gondii* infection and differentiate into ILC1 in the liver [50]. We did not look in the liver for an increase in ILC1, but we did look in the spleen and did not see an increase in CD49a+ ILC1 within the NKp46+ population. We propose that for splenic NK cells and not ILC1 during chronic *T. gondii* infection NKp46 interaction with its ligand and loss of Qa-1b interaction with NKG2A promotes the development of NK cells as negative regulators of CD8+ T cell immunity during chronic *T. gondii* infection.

Our data demonstrates that NK cells present during chronic *T. gondii* infection alter their role in immunity and act as negative regulators of CD8+ T cells to promote reactivation of the parasite. NK cells are the cytotoxic ILC[57]. They also produce high levels of IFNγ and other cytokines after activation. NK cells are usually considered to be a first line of defense against many pathogens and tumors. However, many recent reports demonstrate that NK cells can also negatively regulate adaptive immune responses through several different mechanisms[28; 29; 30; 32; 33; 34; 37; 38; 39; 46; 47; 69]. These include the production of the immunosuppressive cytokine IL-10. NK cells are activated to produce IL-10 during acute stage systemic infections including *T. gondii*. NK cells can also induce apoptosis or kill CD4+ and/or CD8+ T cells during acute infections through TRAIL-TRAILR interactions, NKp46 dependent cytotoxicity and cytotoxicity through undefined receptor ligand pairs. NK cells can also kill tumor infiltrating lymphocytes (TILs) via an NKp46 dependent process. Another study recently published suggests that NK cells that become exhausted during persistent HCV infection lose their ability to produce IFNγ and as a result the CD8+ T cell effector population is unable to be maintained[24]. These studies suggest that NK cells can secrete immune suppressive cytokines to act systemically to suppress immunity, can act directly against T cells and kill them or because they are exhausted themselves they are unable to help maintain CD8+ T cell functions. During chronic *T. gondii* infection we observe that NK cells lose their ability to produce IFNγ while increasing the CD107a expression. Thus while NK cells might lose one function during chronic *T. gondii* infection they appear to have a gain of function. We did attempt to measure whether NK cells from chronically infected mice were more cytotoxic, but we did not observe any increase (data not shown). Importantly CD107a is only a surrogate marker for NK cell cytotoxicity[70]. CD107a can associate with other secretory vesicles and in particular can be surface expressed alongside MHC Class II on DCs[70; 71]. Therefore, we believe that the increase in CD107a on NK cells during chronic *T. gondii* infection may indicate a different type of immune suppressive function. What that suppressive function might be is still unclear. We did not observe NK cells producing IL-10 during the chronic stage of parasite infection and they also did not increase their expression of PDL1, the ligand for PD1. PDL1 expression can promote exhaustion of CD8+ T cells during chronic *T. gondii* infection[10]. Therefore based on our results we propose that NK cells are acquiring a different type of immune suppression than producing IL-10 or being cytotoxic. Another possibility is that sustained NK cell IFNγ is required to help maintain CD8+ T cell function during chronic *T. gondii* infection. NK cells are thought during acute *T. gondii* to help prime CD8+ T cell responses, especially in the absence of CD4+ T cell help[40]. We confirmed the importance of NK cells for priming CD8+ T cells in this study. Based on these studies and our data along with data from the persistent HCV infection study, a lack of NK cell IFNγ could also not support CD8+ T cell function. However, our data show that NK cell depletion enhances CD8+ T cell function during chronic *T. gondii* infection making this less likely and that the negative regulation of CD8+ T cell responses by NK cells is via a different mechanism.

In this study we present data suggesting that during chronic *T. gondii* infection, NK cells are still present, do not appear exhausted based on cell number and PD1 or LAG3 expression. They negatively impact the mortality of chronically infected mice and NK cell depletion rescues animals from CD8+ T cell exhaustion (CD8+ T cell function is maintained and apoptosis reduced) and parasite reactivation. The NK cells develop a unique phenotype and are enriched for cells that are CD49b+ NKp46+ CD94+ NKG2A+ KLRG1+. The development of this population could be dependent upon activating receptor NKp46 recognition of a specific ligand while NKG2A interaction with Qa-1b is reduced. NK cells suppress the CD8+ T cell response by an as of yet identified mechanism that may be independent of cytotoxicity, IL-10 production or the expression of PDL1. Overall in chronic *T. gondii* infection, NK cells may contribute to CD8+ T cell exhaustion and persistence of the parasite and manipulating them to prevent the development of this response could improve health outcomes for individuals susceptible to parasite reactivation.

## Acknowledgements

We would like to acknowledge Corwin Guraedy for animal care and maintenance.

## Author contributions

Ryan Krempels and Jason P. Gigley developed the scientific concept. Ryan Krempels, Daria L. Ivanova, Stephen L. Denton, Kevin D. Fettel, Giandor M. Saltz, David M. Rach, Rida Fatima, Tiffany Mundenke, Joshua Materi and Jason P. Gigley carried out experiments, acquired and analyzed data and helped generate figures for the manuscript. Jason P. Gigley wrote the manuscript. Ildiko R. Dunay consulted on the manuscript.

## Contributions to the field

This study investigated a novel mechanism involved in the development of immune exhaustion during chronic *T. gondii* infection. There are still many unanswered questions about why *T. gondii* is able to persist for life in a host. There are also many open questions about how chronic disease situations cause NK cells to develop responses that can inhibit immunity. Our results demonstrate that in a chronic protozoan infection, NK cells contribute to parasite persistence by enhancing immune exhaustion. The findings also indicate that the chronic inflammatory state of long term *T. gondii* infection modifies the NK cell compartment and that only persistent *T. gondii* infection induces this type of response. This study will help in understanding how to combat life long infection with *T. gondii* to improve therapies for those individuals at high risk for this infection.

## References

[1] P.S. Mead, L. Slutsker, V. Dietz, L.F. McCaig, J.S. Bresee, C. Shapiro, P.M. Griffin, and R.V. Tauxe, Food-related illness and death in the United States. Emerg Infect Dis 5 (1999) 607–25.

[2] J.P. Gigley, The Diverse Role of NK Cells in Immunity to Toxoplasma gondii Infection. PLoS Pathog 12 (2016) e1005396.

[3] G. Harms Pritchard, A.O. Hall, D.A. Christian, S. Wagage, Q. Fang, G. Muallem, B. John, A. Glatman Zaretsky, W.G. Dunn, J. Perrigoue, S.L. Reiner, and C.A. Hunter, Diverse roles for T-bet in the effector responses required for resistance to infection. J Immunol 194 (2015) 1131–40.

[4] I. Coppens, Exploitation of auxotrophies and metabolic defects in Toxoplasma as therapeutic approaches. Int J Parasitol 44 (2014) 109–20.

[5] J.B. Radke, J.N. Burrows, D.E. Goldberg, and L.D. Sibley, Evaluation of Current and Emerging Antimalarial Medicines for Inhibition of Toxoplasma gondii Growth in Vitro. ACS Infect Dis 4 (2018) 1264–1274.

[6] Y. Suzuki, M.A. Orellana, R.D. Schreiber, and J.S. Remington, Interferon-gamma: the major mediator of resistance against Toxoplasma gondii. Science 240 (1988) 516–8.

[7] Y. Suzuki, and J.S. Remington, Dual regulation of resistance against Toxoplasma gondii infection by Lyt-2+ and Lyt-1+, L3T4+ T cells in mice. J Immunol 140 (1988) 3943–6.

[8] R.T. Gazzinelli, S. Hayashi, M. Wysocka, L. Carrera, R. Kuhn, W. Muller, F. Roberge, G. Trinchieri, and A. Sher, Role of IL-12 in the initiation of cell mediated immunity by Toxoplasma gondii and its regulation by IL-10 and nitric oxide. J Eukaryot Microbiol 41 (1994) 9S.

[9] R.T. Gazzinelli, M. Wysocka, S. Hayashi, E.Y. Denkers, S. Hieny, P. Caspar, G. Trinchieri, and A. Sher, Parasite-induced IL-12 stimulates early IFN-gamma synthesis and resistance during acute infection with Toxoplasma gondii. J Immunol 153 (1994) 2533–43.

[10] R. Bhadra, J.P. Gigley, L.M. Weiss, and I.A. Khan, Control of Toxoplasma reactivation by rescue of dysfunctional CD8+ T-cell response via PD-1-PDL-1 blockade. Proc Natl Acad Sci U S A 108 (2011) 9196–201.

[11] R. Bhadra, J.P. Gigley, and I.A. Khan, Cutting edge: CD40-CD40 ligand pathway plays a critical CD8-intrinsic and -extrinsic role during rescue of exhausted CD8 T cells. J Immunol 187 (2011) 4421–5.

[12] R. Bhadra, J.P. Gigley, and I.A. Khan, PD-1-mediated attrition of polyfunctional memory CD8+ T cells in chronic toxoplasma infection. J Infect Dis 206 (2012) 125–34.

[13] S. Hwang, D.A. Cobb, R. Bhadra, B. Youngblood, and I.A. Khan, Blimp-1-mediated CD4 T cell exhaustion causes CD8 T cell dysfunction during chronic toxoplasmosis. J Exp Med (2016).

[14] E.J. Wherry, and M. Kurachi, Molecular and cellular insights into T cell exhaustion. Nat Rev Immunol 15 (2015) 486–99.

[15] T.L. Geiger, and J.C. Sun, Development and maturation of natural killer cells. Curr Opin Immunol 39 (2016) 82–9.

[16] E.Y. Denkers, R.T. Gazzinelli, D. Martin, and A. Sher, Emergence of NK1.1+ cells as effectors of IFN-gamma dependent immunity to Toxoplasma gondii in MHC class I-deficient mice. J Exp Med 178 (1993) 1465–72.

[17] L.L. Johnson, F.P. VanderVegt, and E.A. Havell, Gamma interferon-dependent temporary resistance to acute Toxoplasma gondii infection independent of CD4+ or CD8+ lymphocytes. Infect Immun 61 (1993) 5174–80.

[18] R.T. Gazzinelli, S. Hieny, T.A. Wynn, S. Wolf, and A. Sher, Interleukin 12 is required for the T-lymphocyte-independent induction of interferon gamma by an intracellular parasite and induces resistance in T-cell-deficient hosts. Proc Natl Acad Sci U S A 90 (1993) 6115–9.

[19] C.A. Hunter, C.S. Subauste, V.H. Van Cleave, and J.S. Remington, Production of gamma interferon by natural killer cells from Toxoplasma gondii-infected SCID mice: regulation by interleukin-10, interleukin-12, and tumor necrosis factor alpha. Infect Immun 62 (1994) 2818–24.

[20] J.G. O’Leary, M. Goodarzi, D.L. Drayton, and U.H. von Andrian, T cell- and B cell-independent adaptive immunity mediated by natural killer cells. Nat Immunol 7 (2006) 507–16.

[21] M.A. Cooper, J.M. Elliott, P.A. Keyel, L. Yang, J.A. Carrero, and W.M. Yokoyama, Cytokine-induced memory-like natural killer cells. Proc Natl Acad Sci U S A 106 (2009) 1915–9.

[22] J.C. Sun, J.N. Beilke, and L.L. Lanier, Adaptive immune features of natural killer cells. Nature 457 (2009) 557–61.

[23] S. Paust, H.S. Gill, B.Z. Wang, M.P. Flynn, E.A. Moseman, B. Senman, M. Szczepanik, A. Telenti, P.W. Askenase, R.W. Compans, and U.H. von Andrian, Critical role for the chemokine receptor CXCR6 in NK cell-mediated antigen-specific memory of haptens and viruses. Nat Immunol 11 (2010) 1127–35.

[24] C. Zhang, X.M. Wang, S.R. Li, T. Twelkmeyer, W.H. Wang, S.Y. Zhang, S.F. Wang, J.Z. Chen, X. Jin, Y.Z. Wu, X.W. Chen, S.D. Wang, J.Q. Niu, H.R. Chen, and H. Tang, NKG2A is a NK cell exhaustion checkpoint for HCV persistence. Nat Commun 10 (2019) 1507.

[25] M. Alvarez, F. Simonetta, J. Baker, A. Pierini, A.S. Wenokur, A.R. Morrison, W.J. Murphy, and R.S. Negrin, Regulation of murine NK cell exhaustion through the activation of the DNA damage repair pathway. JCI Insight 5 (2019).

[26] C. Sun, H.Y. Sun, W.H. Xiao, C. Zhang, and Z.G. Tian, Natural killer cell dysfunction in hepatocellular carcinoma and NK cell-based immunotherapy. Acta Pharmacol Sin 36 (2015) 1191–9.

[27] S. Gill, A.E. Vasey, A. De Souza, J. Baker, A.T. Smith, H.E. Kohrt, M. Florek, K.D. Gibbs, Jr., K. Tate, D.S. Ritchie, and R.S. Negrin, Rapid development of exhaustion and down-regulation of eomesodermin limit the antitumor activity of adoptively transferred murine natural killer cells. Blood 119 (2012) 5758–68.

[28] S.N. Waggoner, M. Cornberg, L.K. Selin, and R.M. Welsh, Natural killer cells act as rheostats modulating antiviral T cells. Nature 481 (2012) 394–8.

[29] P.A. Lang, K.S. Lang, H.C. Xu, M. Grusdat, I.A. Parish, M. Recher, A.R. Elford, S. Dhanji, N. Shaabani, C.W. Tran, D. Dissanayake, R. Rahbar, M. Ghazarian, A. Brustle, J. Fine, P. Chen, C.T. Weaver, C. Klose, A. Diefenbach, D. Haussinger, J.R. Carlyle, S.M. Kaech, T.W. Mak, and P.S. Ohashi, Natural killer cell activation enhances immune pathology and promotes chronic infection by limiting CD8+ T-cell immunity. Proc Natl Acad Sci U S A 109 (2012) 1210–5.

[30] K.D. Cook, and J.K. Whitmire, The depletion of NK cells prevents T cell exhaustion to efficiently control disseminating virus infection. J Immunol 190 (2013) 641–9.

[31] D. Peppa, U.S. Gill, G. Reynolds, N.J. Easom, L.J. Pallett, A. Schurich, L. Micco, G. Nebbia, H.D. Singh, D.H. Adams, P.T. Kennedy, and M.K. Maini, Up-regulation of a death receptor renders antiviral T cells susceptible to NK cell-mediated deletion. J Exp Med 210 (2013) 99–114.

[32] J. Crouse, G. Bedenikovic, M. Wiesel, M. Ibberson, I. Xenarios, D. Von Laer, U. Kalinke, E. Vivier, S. Jonjic, and A. Oxenius, Type I interferons protect T cells against NK cell attack mediated by the activating receptor NCR1. Immunity 40 (2014) 961–73.

[33] I.S. Schuster, M.E. Wikstrom, G. Brizard, J.D. Coudert, M.J. Estcourt, M. Manzur, L.A. O’Reilly, M.J. Smyth, J.A. Trapani, G.R. Hill, C.E. Andoniou, and M.A. Degli-Esposti, TRAIL+ NK cells control CD4+ T cell responses during chronic viral infection to limit autoimmunity. Immunity 41 (2014) 646–56.

[34] G. Perona-Wright, K. Mohrs, F.M. Szaba, L.W. Kummer, R. Madan, C.L. Karp, L.L. Johnson, S.T. Smiley, and M. Mohrs, Systemic but not local infections elicit immunosuppressive IL-10 production by natural killer cells. Cell Host Microbe 6 (2009) 503–12.

[35] E. Narni-Mancinelli, B.N. Jaeger, C. Bernat, A. Fenis, S. Kung, A. De Gassart, S. Mahmood, M. Gut, S.C. Heath, J. Estelle, E. Bertosio, F. Vely, L.N. Gastinel, B. Beutler, B. Malissen, M. Malissen, I.G. Gut, E. Vivier, and S. Ugolini, Tuning of natural killer cell reactivity by NKp46 and Helios calibrates T cell responses. Science 335 (2012) 344–8.

[36] D.W. Donley, A.R. Olson, M.F. Raisbeck, J.H. Fox, and J.P. Gigley, Huntingtons Disease Mice Infected with Toxoplasma gondii Demonstrate Early Kynurenine Pathway Activation, Altered CD8+ T-Cell Responses, and Premature Mortality. PLoS One 11 (2016) e0162404.

[37] S.N. Waggoner, K.A. Daniels, and R.M. Welsh, Therapeutic depletion of natural killer cells controls persistent infection. J Virol 88 (2014) 1953–60.

[38] K.D. Cook, H.C. Kline, and J.K. Whitmire, NK cells inhibit humoral immunity by reducing the abundance of CD4+ T follicular helper cells during a chronic virus infection. J Leukoc Biol 98 (2015) 153–62.

[39] C. Rydyznski, K.A. Daniels, E.P. Karmele, T.R. Brooks, S.E. Mahl, M.T. Moran, C. Li, R. Sutiwisesak, R.M. Welsh, and S.N. Waggoner, Generation of cellular immune memory and B-cell immunity is impaired by natural killer cells. Nat Commun 6 (2015) 6375.

[40] C.L. Combe, T.J. Curiel, M.M. Moretto, and I.A. Khan, NK cells help to induce CD8(+)-T-cell immunity against Toxoplasma gondii in the absence of CD4(+) T cells. Infect Immun 73 (2005) 4913–21.

[41] H. Guan, M. Moretto, D.J. Bzik, J. Gigley, and I.A. Khan, NK cells enhance dendritic cell response against parasite antigens via NKG2D pathway. J Immunol 179 (2007) 590–6.

[42] R.S. Goldszmid, A. Bafica, D. Jankovic, C.G. Feng, P. Caspar, R. Winkler-Pickett, G. Trinchieri, and A. Sher, TAP-1 indirectly regulates CD4+ T cell priming in Toxoplasma gondii infection by controlling NK cell IFN-gamma production. J Exp Med 204 (2007) 2591–602.

[43] R.S. Goldszmid, P. Caspar, A. Rivollier, S. White, A. Dzutsev, S. Hieny, B. Kelsall, G. Trinchieri, and A. Sher, NK cell-derived interferon-gamma orchestrates cellular dynamics and the differentiation of monocytes into dendritic cells at the site of infection. Immunity 36 (2012) 1047–59.

[44] D.L. Ivanova, Anti-Asialo GM1 treatment during secondary Toxoplasma gondii infection is lethal and depletes T cells. BioRXIV bioRxiv 550608; doi: (2019).

[45] H.C. Xu, M. Grusdat, A.A. Pandyra, R. Polz, J. Huang, P. Sharma, R. Deenen, K. Kohrer, R. Rahbar, A. Diefenbach, K. Gibbert, M. Lohning, L. Hocker, Z. Waibler, D. Haussinger, T.W. Mak, P.S. Ohashi, K.S. Lang, and P.A. Lang, Type I interferon protects antiviral CD8+ T cells from NK cell cytotoxicity. Immunity 40 (2014) 949–60.

[46] S.Q. Crome, L.T. Nguyen, S. Lopez-Verges, S.Y. Yang, B. Martin, J.Y. Yam, D.J. Johnson, J. Nie, M. Pniak, P.H. Yen, A. Milea, R. Sowamber, S.R. Katz, M.Q. Bernardini, B.A. Clarke, P.A. Shaw, P.A. Lang, H.K. Berman, T.J. Pugh, L.L. Lanier, and P.S. Ohashi, A distinct innate lymphoid cell population regulates tumor-associated T cells. Nat Med 23 (2017) 368–375.

[47] B. Kwong, R. Rua, Y. Gao, J. Flickinger, Jr., Y. Wang, M.J. Kruhlak, J. Zhu, E. Vivier, D.B. McGavern, and V. Lazarevic, T-bet-dependent NKp46+ innate lymphoid cells regulate the onset of TH17-induced neuroinflammation. Nat Immunol 18 (2017) 1117–1127.

[48] L.L. Lanier, NK cell recognition. Annu Rev Immunol 23 (2005) 225–74.

[49] D.L. Ivanova, R. Fatima, and J.P. Gigley, Comparative Analysis of Conventional Natural Killer Cell Responses to Acute Infection with Toxoplasma gondii Strains of Different Virulence. Front Immunol 7 (2016) 347.

[50] E. Park, S. Patel, Q. Wang, P. Andhey, K. Zaitsev, S. Porter, M. Hershey, M. Bern, B. Plougastel-Douglas, P. Collins, M. Colonna, K.M. Murphy, E. Oltz, M. Artyomov, L.D. Sibley, and W.M. Yokoyama, Toxoplasma gondii infection drives conversion of NK cells into ILC1-like cells. Elife 8 (2019).

[51] A. Diefenbach, M. Colonna, and S. Koyasu, Development, differentiation, and diversity of innate lymphoid cells. Immunity 41 (2014) 354–365.

[52] G. Eberl, J.P. Di Santo, and E. Vivier, The brave new world of innate lymphoid cells. Nat Immunol 16 (2015) 1–5.

[53] C.S.N. Klose, M. Flach, L. Mohle, L. Rogell, T. Hoyler, K. Ebert, C. Fabiunke, D. Pfeifer, V. Sexl, D. Fonseca-Pereira, R.G. Domingues, H. Veiga-Fernandes, S.J. Arnold, M. Busslinger, I.R. Dunay, Y. Tanriver, and A. Diefenbach, Differentiation of type 1 ILCs from a common progenitor to all helper-like innate lymphoid cell lineages. Cell 157 (2014) 340–356.

[54] C.A. Hunter, R. Chizzonite, and J.S. Remington, IL-1 beta is required for IL-12 to induce production of IFN-gamma by NK cells. A role for IL-1 beta in the T cell-independent mechanism of resistance against intracellular pathogens. J Immunol 155 (1995) 4347–54.

[55] D.L. Ivanova, T.M. Mundhenke, and J.P. Gigley, The IL-12- and IL-23-Dependent NK Cell Response Is Essential for Protective Immunity against Secondary Toxoplasma gondii Infection. J Immunol 203 (2019) 2944–2958.

[56] S. Kim, J. Poursine-Laurent, S.M. Truscott, L. Lybarger, Y.J. Song, L. Yang, A.R. French, J.B. Sunwoo, S. Lemieux, T.H. Hansen, and W.M. Yokoyama, Licensing of natural killer cells by host major histocompatibility complex class I molecules. Nature 436 (2005) 709–13.

[57] V.S. Cortez, and M. Colonna, Diversity and function of group 1 innate lymphoid cells. Immunol Lett 179 (2016) 19–24.

[58] A. Vitenshtein, Y. Charpak-Amikam, R. Yamin, Y. Bauman, B. Isaacson, N. Stein, O. Berhani, L. Dassa, M. Gamliel, C. Gur, A. Glasner, C. Gomez, R. Ben-Ami, N. Osherov, B.P. Cormack, and O. Mandelboim, NK Cell Recognition of Candida glabrata through Binding of NKp46 and NCR1 to Fungal Ligands Epa1, Epa6, and Epa7. Cell Host Microbe 20 (2016) 527–534.

[59] O. Mandelboim, N. Lieberman, M. Lev, L. Paul, T.I. Arnon, Y. Bushkin, D.M. Davis, J.L. Strominger, J.W. Yewdell, and A. Porgador, Recognition of haemagglutinins on virus-infected cells by NKp46 activates lysis by human NK cells. Nature 409 (2001) 1055–60.

[60] Y. Bar-On, Y. Charpak-Amikam, A. Glasner, B. Isaacson, A. Duev-Cohen, P. Tsukerman, A. Varvak, M. Mandelboim, and O. Mandelboim, NKp46 Recognizes the Sigma1 Protein of Reovirus: Implications for Reovirus-Based Cancer Therapy. J Virol 91 (2017).

[61] S.H. Lee, K.S. Kim, N. Fodil-Cornu, S.M. Vidal, and C.A. Biron, Activating receptors promote NK cell expansion for maintenance, IL-10 production, and CD8 T cell regulation during viral infection. J Exp Med 206 (2009) 2235–51.

[62] T.A. Holderried, P.A. Lang, H.J. Kim, and H. Cantor, Genetic disruption of CD8+ Treg activity enhances the immune response to viral infection. Proc Natl Acad Sci U S A 110 (2013) 21089–94.

[63] R.E. Vance, J.R. Kraft, J.D. Altman, P.E. Jensen, and D.H. Raulet, Mouse CD94/NKG2A is a natural killer cell receptor for the nonclassical major histocompatibility complex (MHC) class I molecule Qa-1(b). J Exp Med 188 (1998) 1841–8.

[64] H. Spits, D. Artis, M. Colonna, A. Diefenbach, J.P. Di Santo, G. Eberl, S. Koyasu, R.M. Locksley, A.N. McKenzie, R.E. Mebius, F. Powrie, and E. Vivier, Innate lymphoid cells--a proposal for uniform nomenclature. Nat Rev Immunol 13 (2013) 145–9.

[65] J.C. Sun, Transcriptional Control of NK Cells. Curr Top Microbiol Immunol 395 (2016) 1–36.

[66] P.H. Kruse, J. Matta, S. Ugolini, and E. Vivier, Natural cytotoxicity receptors and their ligands. Immunol Cell Biol 92 (2014) 221–9.

[67] H.J. Pegram, D.M. Andrews, M.J. Smyth, P.K. Darcy, and M.H. Kershaw, Activating and inhibitory receptors of natural killer cells. Immunol Cell Biol 89 (2011) 216–24.

[68] M.S. Tessmer, C. Fugere, F. Stevenaert, O.V. Naidenko, H.J. Chong, G. Leclercq, and L. Brossay, KLRG1 binds cadherins and preferentially associates with SHIP-1. Int Immunol 19 (2007) 391–400.

[69] J. Crouse, H.C. Xu, P.A. Lang, and A. Oxenius, NK cells regulating T cell responses: mechanisms and outcome. Trends Immunol 36 (2015) 49–58.

[70] G. Alter, J.M. Malenfant, and M. Altfeld, CD107a as a functional marker for the identification of natural killer cell activity. J Immunol Methods 294 (2004) 15–22.

[71] X. Michelet, S. Garg, B.J. Wolf, A. Tuli, P. Ricciardi-Castagnoli, and M.B. Brenner, MHC class II presentation is controlled by the lysosomal small GTPase, Arl8b. J Immunol 194 (2015) 2079–88.

